# Epitope Mapping of SARS-CoV-2 Spike Protein Reveals Distinct Antibody Binding Activity of Vaccinated and Infected Individuals

**DOI:** 10.1101/2022.04.13.487697

**Authors:** Nathaniel Felbinger, David Trudil, Lawrence Loomis, Richard Ascione, Gregory Siragusa, Seiji Haba, Shruti Rastogi, Aidan Mucci, Mark Claycomb, Sebastian Snowberger, Brian Luke, Stephen Francesconi, Shirley Tsang

**Affiliations:** New Horizons Diagnostics, Inc., Arbutus, MD; Georgetown University School of Medicine, Washington, DC; Scout Microbiology LLC, Waukesha, WI; Frederick, MD 21702; DTRA (Defense Threat Reduction Agency), Ft. Belvoir, VA

**Keywords:** B-cell epitope, variant mutations, peptide microarray, IgG, covid spike protein

## Abstract

Previous studies have attempted to characterize the antibody response of individuals to the SARS-CoV-2 virus on a linear peptide level by utilizing peptide microarrays. These studies have helped to identify epitopes that have potential to be used for diagnostic tests to identify infected individuals, however, the immunological responses of individuals who have received the currently available Moderna mRNA-1273 or Pfizer BNT162b2 mRNA vaccines have not been characterized. We aimed to identify linear peptides of the SARS-CoV-2 spike protein that elicited high IgG or IgA binding activity and to compare the immunoreactivity of infected individuals to those who received both doses of either vaccines by utilizing peptide microarrays. Our results revealed peptide epitopes of significant IgG binding among recently infected individuals. Some of these peptides are located near functional domains implicated in the high infectivity of SARS-CoV-2. Vaccinated individuals lacked these distinct markers despite overall binding activity being similar.

## INTRODUCTION

Emerging variants of SARS-CoV-2 continue to make up an increased proportion of reported cases (1). These variants contain mutations that play a role in viral infectivity and decrease the effectiveness of the current vaccines (2, 3), increasing the importance of epitope-based studies of the antibody response of vaccinated individuals and previously infected individuals. Studying epitopes at the peptide level helps to reveal which mutations in currently prevalent variants may increase infectivity and transmissibility. In addition, epitopes that are found to have consistent antibody binding may serve as a useful diagnostic target to assess previous infection or the general protective status of individuals. Viral load detected using PCR testing can fall to undetectable levels within one to two weeks after symptom onset (4) and antigen tests are sometimes plagued by low sensitivity (5). Gaining an understanding of the epitopes that elicit significant antibody binding may prove exceptionally useful for improving diagnostic methods and vaccine development.

Peptide microarrays are a tool with many immunological applications, including the screening of antibody binding activity against antigens or entire pathogenic proteomes. This information can provide answers regarding the presence of potential epitopes and characterization of the target pathogen’s mechanisms of infection. As entire targeted viral proteomes can be bound to microarrays, the process of screening for antibody binding is both fast and reproducible. These capabilities alongside streamlined data outputs make the peptide microarray a particularly powerful tool to perform systematic epitope profiling of viral pathogens such as SARS-CoV-2.

The convenient systematic approach to assessing individual peptides as potential epitopes using peptide microarrays may prove especially important with the global relevance of the Omicron variant which arose in the winter of 2021 (6). The Omicron variant displays apparent lowered pathogenicity (7) while at the same time maintaining higher virulence, helping it to become the current dominant strain of SARS-CoV-2. This extreme virulence has coincided with the rise of Omicron subvariants (8), which appear posed to continue increasing in number. Proper assessment of the mutations found in the subvariants is necessary to determine their effect on the virulence and pathogenicity of the virus.

Previous works utilizing peptide microarrays have focused heavily on mapping the humoral responses to the SARS-CoV-2 proteome using samples from individuals currently or recently infected with SARS-CoV-2 (9–15). SARS-CoV-2 epitopes that elicit a significant IgG and/or IgM antibody response in infected or previously infected samples have been identified in the spike (S), nucleocapsid (N), membrane (M), Orf1ab, and Orf3a proteins. While cross-reactivity with other common coronaviruses has been observed, (5,7) there are many epitopes that demonstrate highly specific reactivity with SARS-CoV-2 antibodies distributed across the SARS-CoV-2 proteome. Many epitopes have been identified in previous studies (11–13), both in commercially available and independently designed peptide microarrays. These findings support the use of the peptide microarray approach in locating highly specific and sensitive SARS-CoV-2 human B Cell epitopes.

Several studies (9–15) have demonstrated that peptide microarrays can discriminate between mild symptomatic, severe symptomatic, and eventually fatal cases of SARS-CoV-2 based on the antibody reactivity of specific SARS-CoV-2 epitopes (12, 15). Additional microarray studies have also demonstrated that specific epitopes identified using a peptide microarray can elicit a strong neutralizing antibody response (15). These findings indicate that there may exist a linear epitope or group of linear epitopes that can serve as a marker of the protective status of individuals against SARS-CoV-2 infection. While this is an exciting prospect, previous microarray epitope mapping studies have been focused exclusively on the comparison of infected and naive individuals. To our knowledge no study has utilized a peptide-microarray approach to study individuals who received the two currently available mRNA vaccines, Moderna mRNA-1273 and Pfizer BNT162b2. It is to be expected that a peptide microarray is capable of differentiating the immune response of vaccinated individuals by analyzing the antibody binding to the targeted SARS-CoV-2 spike protein peptides.

Herein we report a study of IgG and IgA antibody reactivity of individuals vaccinated with the Moderna or Pfizer vaccines against linear peptides. As the Moderna and Pfizer vaccines contain the sequences for the SARS-CoV-2 spike protein, our study has focused mainly on the spike protein region. Data herein identifies epitopes that have the potential to serve as markers to assess SARS-CoV-2 protection in addition to gaining a general characterization of IgG and IgA antibody response against SARS-CoV-2 in vaccinated individuals. An assessment of the potential impact of mutations found in current prevalent variants within these identified epitopes was also performed. These epitopes may be useful in the study of immune response development in infected and vaccinated individuals and aid in the development of immunity or detection assays.

## MATERIALS AND METHODS

### 1.1 Infected, Vaccinated, and Control Serum Samples

Serum samples were purchased from RayBiotech (Atlanta, GA) and Reprocell (Beltsville, MD). Samples were divided into three groups:

- SARS-Cov-2 positive individuals (n=30)
- Individuals who received either the Moderna or Pfizer mRNA vaccines (n=17)
- A naive negative control group (n=10)

Among the 10 negative controls, five serum samples were collected prior to the pandemic before the end of 2019 (n=5). There are five paired vaccinated individuals whose serum samples were collected one to three days before receiving the first dose of either the Moderna (n=2) or Pfizer (n=3) vaccine. These five were reported as never having been infected before their vaccination. The post-vaccinated sera were collected between 6 - 44 days after receiving their second dose of either Moderna or Pfizer vaccine. Basic demographic information and date of sample collection relative to symptom onset for infected patients can be found in Supplemental Table S1. Demographic information for vaccinated and negative subjects can be found in Supplemental Table S2. Available symptom information for all infected samples is available in Supplemental Table S3. All purchased infected sera was inactivated with a 4.0% Triton X-100 treatment prior to their arrival at our facility. All samples were immediately aliquoted and stored at −80° C upon arrival to our facility.

### 1.2 Peptide Microarrays

For our systematic proteome analysis, PEPperCHIP® (Heidelberg, Germany) SARS-CoV-2 Proteome Microarray slides were used for all samples. These slides contain a series of linear peptide sequences that cover the entire SARS-COV-2 proteome. Each of these peptides are printed in duplicate and consist of 15 amino acids in total, 13 of which overlap with neighboring SARS-CoV-2 peptide sequences. Additionally, the slides contain certain mutated peptide sequences found in SARS-CoV-2 variants and a set of influenza and polio peptides which were utilized as positive controls.

Prior to performing the procedure, solutions of PBST wash buffer (Phosphate-buffered saline with 0.05% Tween20, at 7.4 pH), and pH 7.4 1mM Tris dipping buffer were prepared. All solutions were filtered using a 0.44-micron vacuum filter kit and pH was adjusted to 7.4 with the addition of 3M HCl if necessary. Microarray slides were initially incubated in a solution of 0.05% PBST (7.4 pH) and 10% blocking buffer (Rockland Immunochemicals, Inc., Pottstown, PA) for 15 minutes at 23°C. Microarray slides were aspirated and then blocked with blocking buffer for 30 minutes at 23°C. After blocking, slides were again aspirated and probed with a mixture of fluorescent secondary antibodies that would be used for peptide binding detection consisting of Rabbit Anti-Human IgG DyLight™ 800 (Rockland Immunochemicals), Alexa Conjugated Goat Anti-Human IgA 680 (Jackson Immuno Research Labs, West Grove, PA), and Mouse Anti-HA (influenza virus hemagglutinin) Dylight™ 680 (PEPperCHIP®). This prescreen of secondary antibody treatment was used to reveal any non-specific secondary antibody interactions with the slide.

After performing three PBST washes of one minute each, slides were treated with sera diluted 1:500 in a 10% blocking buffer/PBST solution and incubated overnight at 4°C. Following this overnight incubation, slides were washed three times with PBST for one minute each and dipped into the previously prepared 1mM Tris Solution. Detection of IgG was performed via treatment with Rabbit Anti-Human IgG DyLight™ 800 for 45 minutes at 23°C. After performing three PBST washes of one minute each, slides were then treated a second time with samples diluted 1:200 in 10% blocking buffer/ PBST and incubated overnight at 4°C. After the second overnight incubation, samples were washed three times with PBST for one minute each and treated with Alexa Conjugated Goat Anti-Human IgA 680 for 45 minutes at 23°C for detection of IgA. Before scanning, a final set of three one-minute washes with PBST was performed followed by dipping the slides three times in the 1mM Tris Solution.

Fluorescent signals were acquired using the InnoScan 710-IR microarray scanner from Innopsys (Chicago, IL) (4). Slides were scanned at wavelengths of 670 nm for IgA detection and 785 nm for IgG detection and at a resolution of 30uM for both wavelengths. Fluorescent values were retrieved using Innopsys Mapix Microarray Analysis Software.

### 1.3 Epitope Validation

The tiff image files, acquired after probing slides with secondary antibody solution, were analyzed for any slide artifacts prior to data analysis (16). Artifact sequences from the prescreen of secondary antibody treatment were excluded from further analysis. Any microarray slides that did not display sufficient reactivity with polio and HA positive control peptide spots were considered invalid.

For data analysis, we used the background-subtracted median intensity values acquired from the Mapix Microarray Image Acquisition and Analysis Software (Innopsys). To identify potential epitopes, we first averaged these median fluorescent intensity values of each peptide found within the spike protein for each experimental group. We then selected peptides in which the average median intensity value of the infected group was at least 1.5-fold greater and had p-values of < 0.05 from an unpaired t-test comparison to the average log_2_-normalized median intensity value of the negative controls. Among these selected peptides, we compared the average log_2_-normalized median intensity value of each peptide to their neighboring peptides. Peptides that had significantly different fluorescent intensity values were screened out, and the remaining peptides were recognized as potential epitopes (5, 17)

### 1.4 ELISA Testing

In-house ELISA tests were utilized to detect levels of IgG and IgA antibodies against SARS-CoV-2 receptor-binding domain (RBD) antigen. Samples were diluted at 1:10,000 in a 1% milk PBST solution. 100 μl of serum samples (in duplicate) were added to ELISA plates that were coated with 150 ng of SARS-CoV-2 RBD antigen per well. The plates were incubated overnight at 4°C and washed three times in a 0.1% PBST solution. Secondary detection antibody solutions of Mouse Anti-Human IgG Fc-HRP (SouthernBiotech, Birmingham, AL) diluted 1:10,000 in 1% milk PBST and Mouse Anti-Human IgA1-HRP/Mouse Anti-Human IgA2-HRP (SouthernBiotech) diluted 1:4,000 in 1% milk PBST solution were used to detect IgG and IgA, respectively. Secondary antibody solutions were incubated at room temperature for one hour on an orbital shaker set at 200 rpm. Plates were washed three times with a 1% PBST solution and developed for 10 minutes with a TMB substrate solution. 3M HCl was added and plates were scanned using the SpectraMax® iD3 Multi-Mode Microplate Reader (Molecular Devices, San Jose, CA) (Table S4).

## RESULTS

### 2.1 SARS-CoV-2 Spike Epitopes Exhibiting Significant IgG Binding Identified in Infected Samples

To identify reactive epitopes, we looked at peptides in which the average log_2_-normalized intensity value of either of the two infected sample groups had a fold-change value of at least 1.5 and yielded adjusted p-values of < 0.05 from an unpaired t-test when compared to the mean log_2_-normalized intensity value from the negative control group (15, 17). Using these criteria, nine peptides with significantly higher IgG antibody binding were identified in the spike protein region (Figure 1a and Table 1) among infected individuals.

**Figure 1a.**
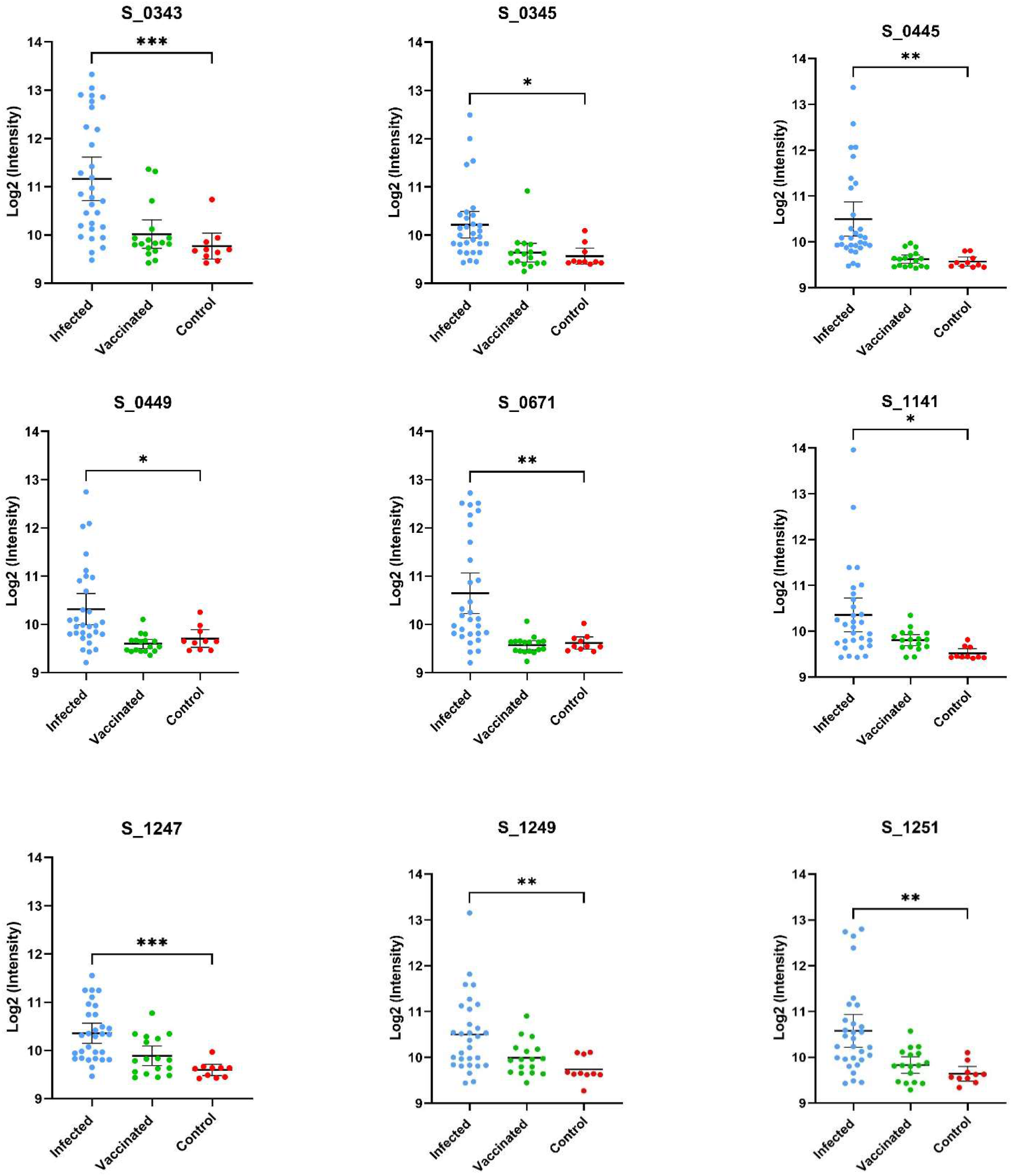
Displayed are the log_2_ normalized individual fluorescent intensity values of infected (n=30), vaccinated (n=17), and control (n=10) sample groups. The peptide ID along with the results of the unpaired t-test comparing the infected and control groups can be found above the data. The results of the t-test comparing the fluorescent intensities of the infected and negative control groups values are shown (* < .05, ** < .01, *** < .001, **** < .0001).

**Table 1.**
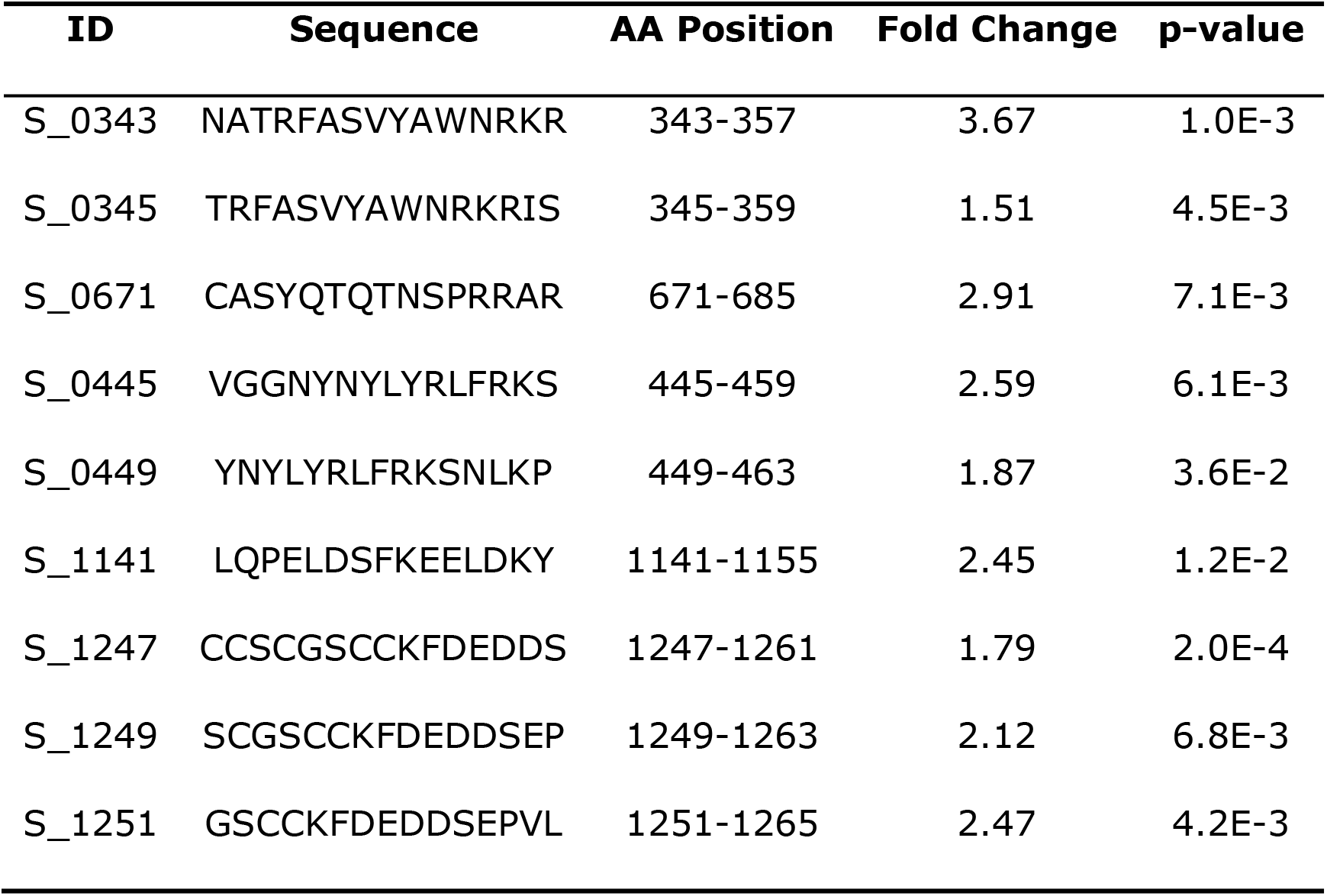
Listed are the peptides that met our criteria to be classified as potential epitopes among infected individuals. The leftmost column contains peptide IDs, which were assigned based on the position of the first amino acid in the sequence. Amino acid sequences and their location in the spike protein sequence are displayed in the second and third columns, respectively. Fold change and p-values acquired are displayed in the fourth and fifth columns respectively

Using the same approach of epitope identification with our vaccinated groups, no peptides were found to meet the criteria for significant IgG binding activity. Two peptides were found to have p-values less than 0.05, but these peptides did not meet our fold change criteria as described in our methods (Figure 1b). Of the nine peptides identified as significant in the infected sample group, five were also found to have significantly greater IgG binding than the vaccine group. Peptides with no significant differences in binding activity between the recently infected group and vaccinated group were peptides S_1141 and S_1247-S_1251. The SARS-CoV-2 RBD IgG titers of the vaccinated individuals were comparable to the IgG titers of infected samples according to data supplied by the sample provider, RayBiotech, and our own in-house RBD ELISA testing (Table S4).

**Figure 1b.**
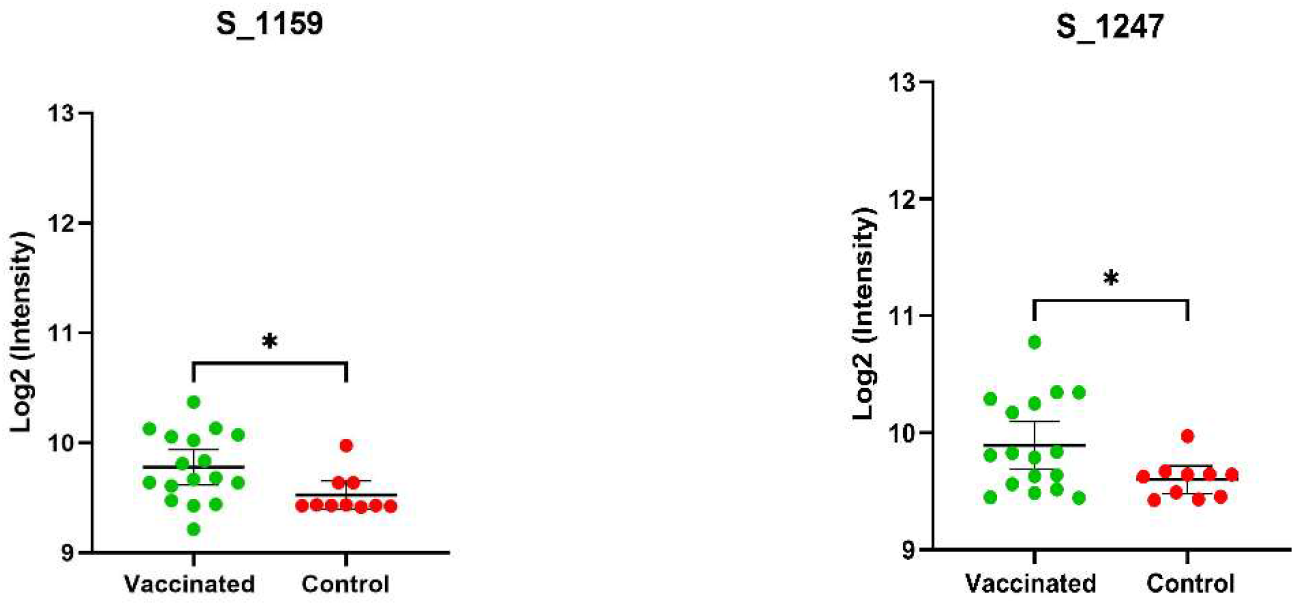
Depicted above are mean log_2_ normalized fluorescent intensities of the vaccinated and control groups that were found to be significantly different (p-value < .05) based on the results of an unpaired t-test. The ID and results of the t-test are displayed above the plots. The vaccinated group was not found to have a fold change < 1.5 and, therefore, these peptides were not recognized as epitopes of highest interest based on our criteria.

Within the infected samples, peptides identified as having significant IgG binding activity appear to be distributed throughout the spike protein (Figure 2). Identified reactive epitopes that meet our criteria include two consecutive peptides in the RBD (S_0343 and S_0345), one located in the putative fusion peptide domain (S_0671), one found in proximity to the identified Heptad Repeat 2 sequence (S_1141), and three located within a cysteine-rich sequence (S_1247, S_1249, and S_1251) neighboring what is thought to be a transmembrane domain (18) located in the C terminal of the S2 subunit. Among these peptides, those that were found to have relatively higher intensity among the vaccinated group (S_1159 and S_1247) were both found in the S2 region. Seven frequent mutations in common variants were identified within at least one peptide sequence (Table 3) and were found to be well distributed throughout the spike protein.

**Figure 2.**
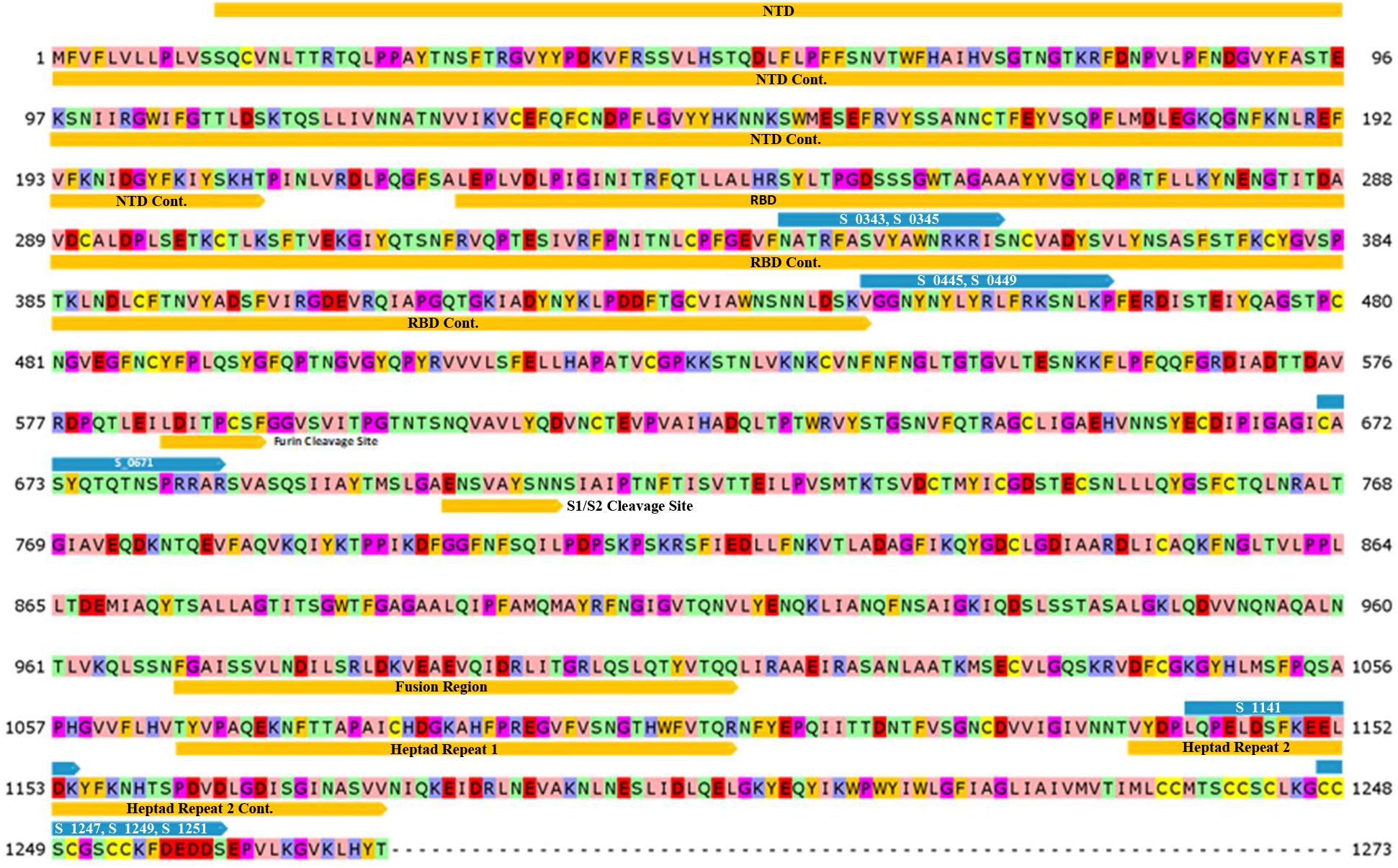
The protein sequence of the SARS-CoV-2 spike protein was annotated to study the proximity of identified potential epitopes to functional regions. Blue bars depict the locations of epitopes with significant IgG binding. Orange bars cover regions that are associated with predicted protein function. Protein sequence is colored according to the Zappo amino acid color scheme (with brightness of some colors increased for clarity).

### 2.2 No Significant Difference in IgG Epitope Profiles of Individuals who Received the Moderna or Pfizer Vaccine

No individual linear peptide exhibited significant IgG binding in our vaccinated sample group as per our criteria. Overall IgG binding activity in the vaccinated group was relatively comparable to the infected group. However, no individual peptide with consistent binding emerged in the vaccinated group as significant. To ascertain differences in peptide epitope binding activity, we compared the IgG binding of individuals who received the Moderna (n=8) and the Pfizer mRNA vaccines (n=9). Comparison of the Moderna and Pfizer vaccinated sample groups revealed little in terms of significant differences in IgG epitope binding activity in samples between the two vaccines. Both vaccinated groups appeared to have more IgG binding activity to epitopes in the S2 region of the spike protein (Figure 3), although peptide S_0343 located in the RBD region appeared to have significant binding in certain individuals of both groups. To compare the peptides that elicited the highest IgG binding in both groups, we identified peptides that displayed fluorescent intensity values that were 3.0 standard deviations above the mean in each group (Table 2). Eleven peptides were found to have intensities above this cutoff in the Moderna sample group and two peptides were identified as being above this cutoff in the Pfizer sample group. Both peptides above 3.0 deviations in the Pfizer vaccinated group were also found among the peptides identified in the Moderna vaccinated group. Of the eleven peptides identified in the Moderna vaccinated sample group, six were either identified as having significant IgG binding activity or overlapped with a peptide found to have significant IgG binding activity within the infected group.

**Figure 3.**
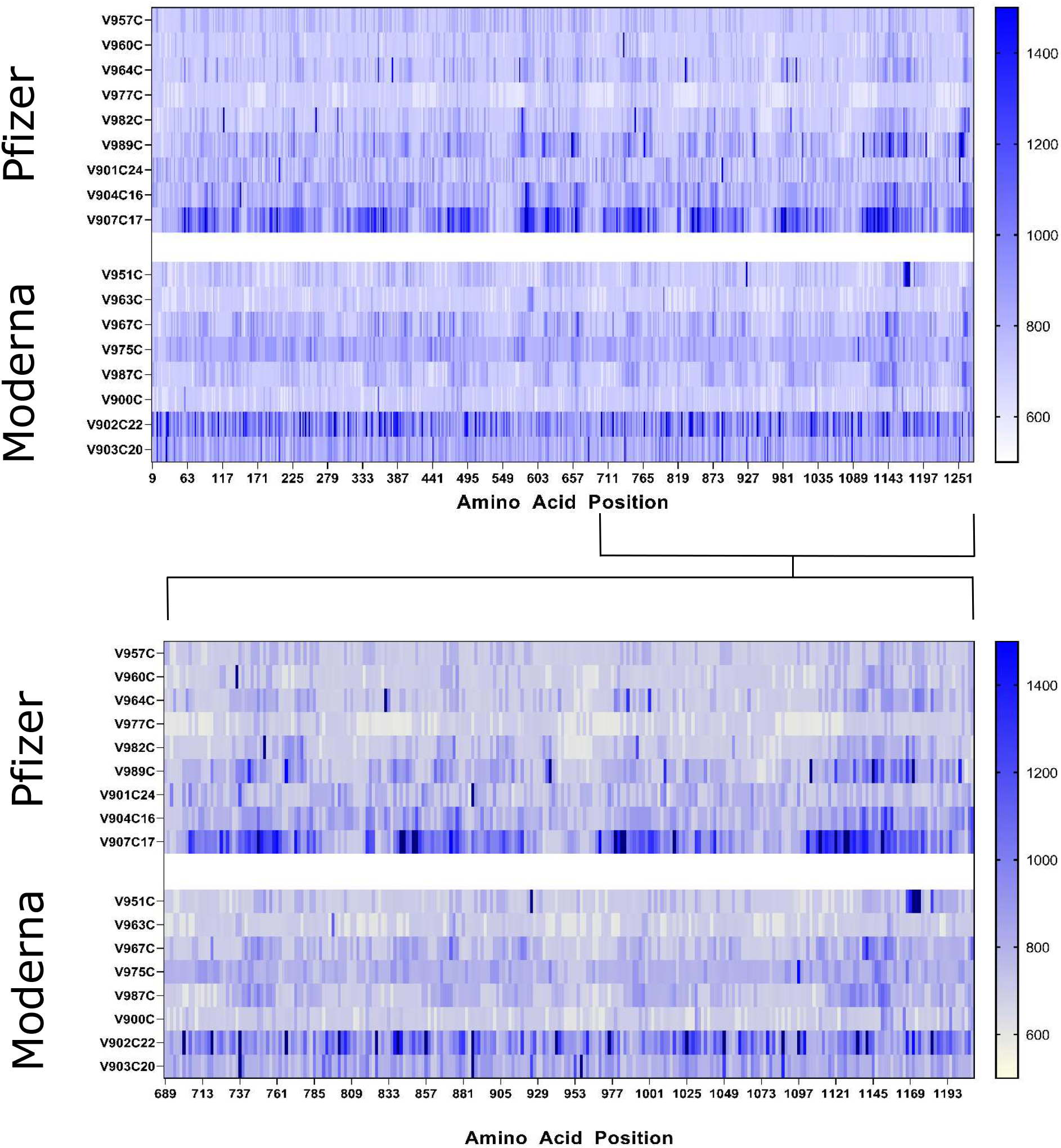
Heat maps of the whole SARS-CoV-2 spike protein (top) and S2 region (bottom) displaying the raw fluorescent intensity values of vaccinated samples were created to compare epitope binding of Moderna and Pfizer vaccinated individuals. The fluorescent intensity values were read at the 785 nm wavelength, which corresponds to IgG binding activity. Sample IDs are displayed to the left of their respective rows. Individuals who received the Pfizer vaccine are displayed in the upper panels while those who received the Moderna vaccine are found in the lower panels. Amino acid sequence of the entire spike protein is displayed in a linear manner with amino acid positions denoted below.

**Table 2.**
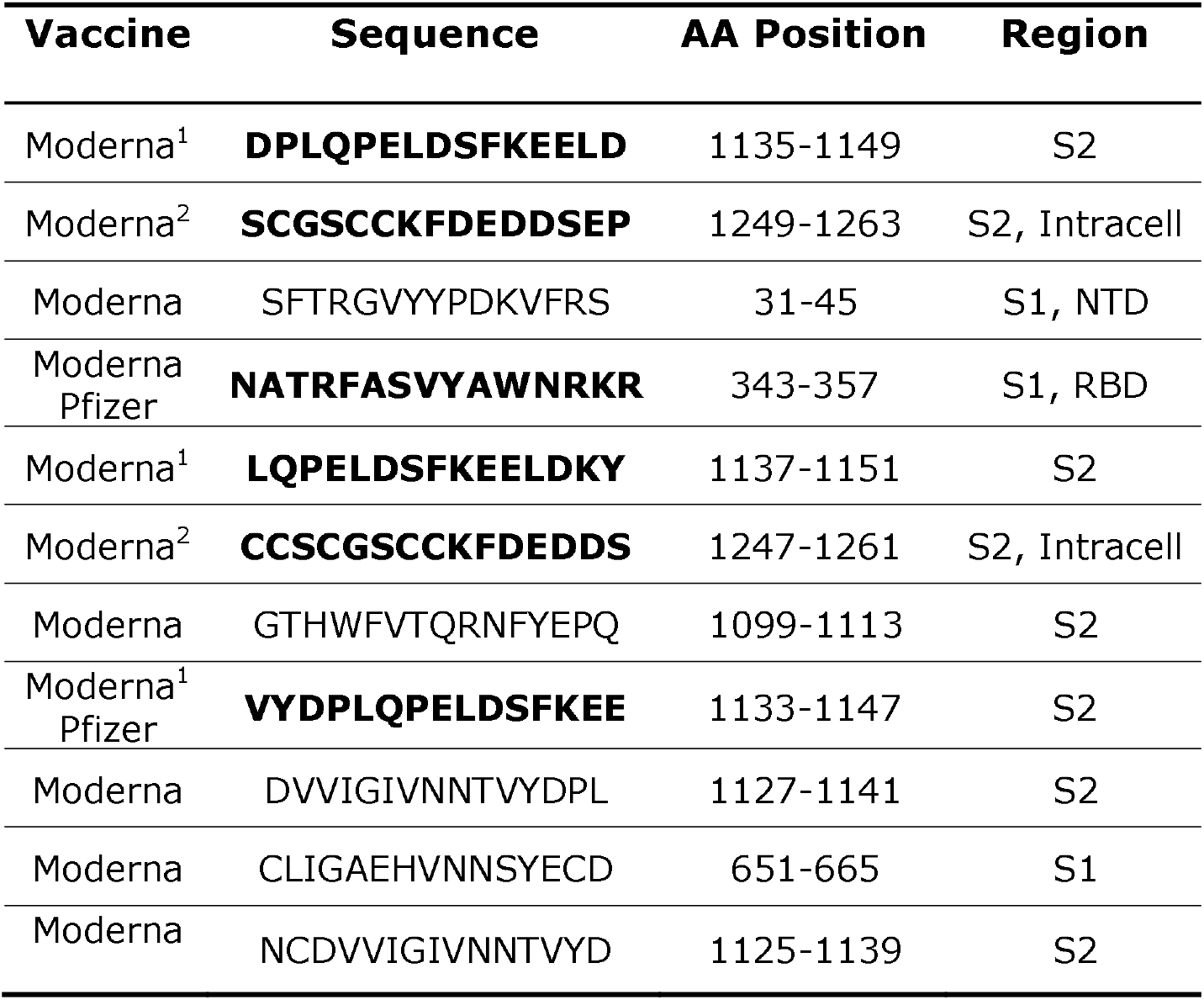
Presented are peptides whose mean fluorescent intensities were at least three standard deviations higher than the overall average intensities in either the Moderna or Pfizer vaccinated groups. The vaccine group in which the peptide met these criteria are displayed in the first column (no peptides were found to be unique to the individuals who received the Pfizer vaccine). Peptide sequences are displayed in the second column. Bolded sequences are among or share significant amino acid identity with one of the 10 potential epitopes identified in the infected sample group. Amino acid position and domain/subregion of each sequence is displayed in the third and fourth columns. 1 Part of the contiguous amino acid sequence VYDPLQPELDSFKEELDKY. 2 Part of the contiguous amino acid sequence CCSCGSCCKFDEDDSEP.

### 2.3 No Significant IgA Binding Activity to SARS-CoV-2 Spike Protein Epitopes Identified in Infected or Vaccinated Individuals

The same approach used to identify IgG epitopes was utilized to identify IgA epitopes. However, no significant IgA epitopes were identified in either the infected group or the vaccinated group. The IgA titer information of the samples provided by RayBiotech was limited, but in-house ELISA testing of the RBD antigen revealed low levels of IgA binding throughout both infected and vaccinated groups (Table S4). The PEPperCHIP® microarrays contained the entire proteome of SARS-CoV-2, allowing for identification of significant IgA binding to peptides outside of the spike protein. As the scope of this report is solely focused on the spike protein these epitopes have been excluded.

## DISCUSSION

To assess the potential of incorporating the discovered peptide sequences into viable therapeutics or diagnostics, we investigated the conservation of sequences among other closely related coronaviruses and the proximity of the sequences to mutations found in current dominant variants. Previous studies have established that antibodies that bind to certain linear epitopes found in the SARS-CoV-2 proteome are found to be reactive to conserved peptide stretches in other coronaviruses (CoVs) (6). Aligning the epitopes to closely related CoVs helped to assess the potential for nonspecific antibody binding by antibodies present due to exposure to other coronaviruses. With the current global prevalence of variant strains such as Delta and Omicron, it is important to investigate if the identified peptide sequences will be affected by common mutations of the spike protein.

### 3.1 Alignment Analysis of Identified Potential Peptide Epitopes

There are currently six identified human infecting coronaviruses (hCoVs) other than SARS-CoV-2. Four of these coronaviruses, the alphacoronaviruses HCoV-229E and HCoV-NL63 and betacoronaviruses HCoV-OC43 (lineage A) and HCoV-HKU1 (lineage A) are highly virulent and associated with causing upper respiratory illness in human adults (19). The other two hCoVs are SARS-CoV (lineage B) and MERS-CoV (lineage C), which are less virulent and more pathogenic. To predict potential antibodies that may exhibit cross-reactive binding with peptides of other CoVs, we created alignments of the spike protein sequences found in the seven hCoVs as well as the bat-infecting CoV RaTG13 (lineage B) and Pangolin-CoV (lineage B) (Figure 4). The alignments revealed that the identified peptides were poorly conserved outside of the lineage B hCoVs. Of the nine peptides identified among the acutely infected group, the peptides S_1141 and S_1247-S_1251 were found to be the most conserved among the hCoVs. Peptide S_1141, located in proximity to the HR2 (heptad repeat domain 2) sequence, was found to have a relatively similar identity with 6/15 amino acid residues being shared among all the peptides and four of the other amino acid residues being classified as conserved or semi-conserved sequence alterations. The sequence in proximity to S_1141 has also been identified as being potentially cross-reactive in previous microarray studies of SARS-CoV-2 (12). Peptides S_1247-S_1251 are located near the C-terminus of the spike protein just outside the transmembrane domain. These contiguous peptides contain highly conserved cysteine residues which are thought to be imperative for membrane fusion (20).

**Figure 4.**
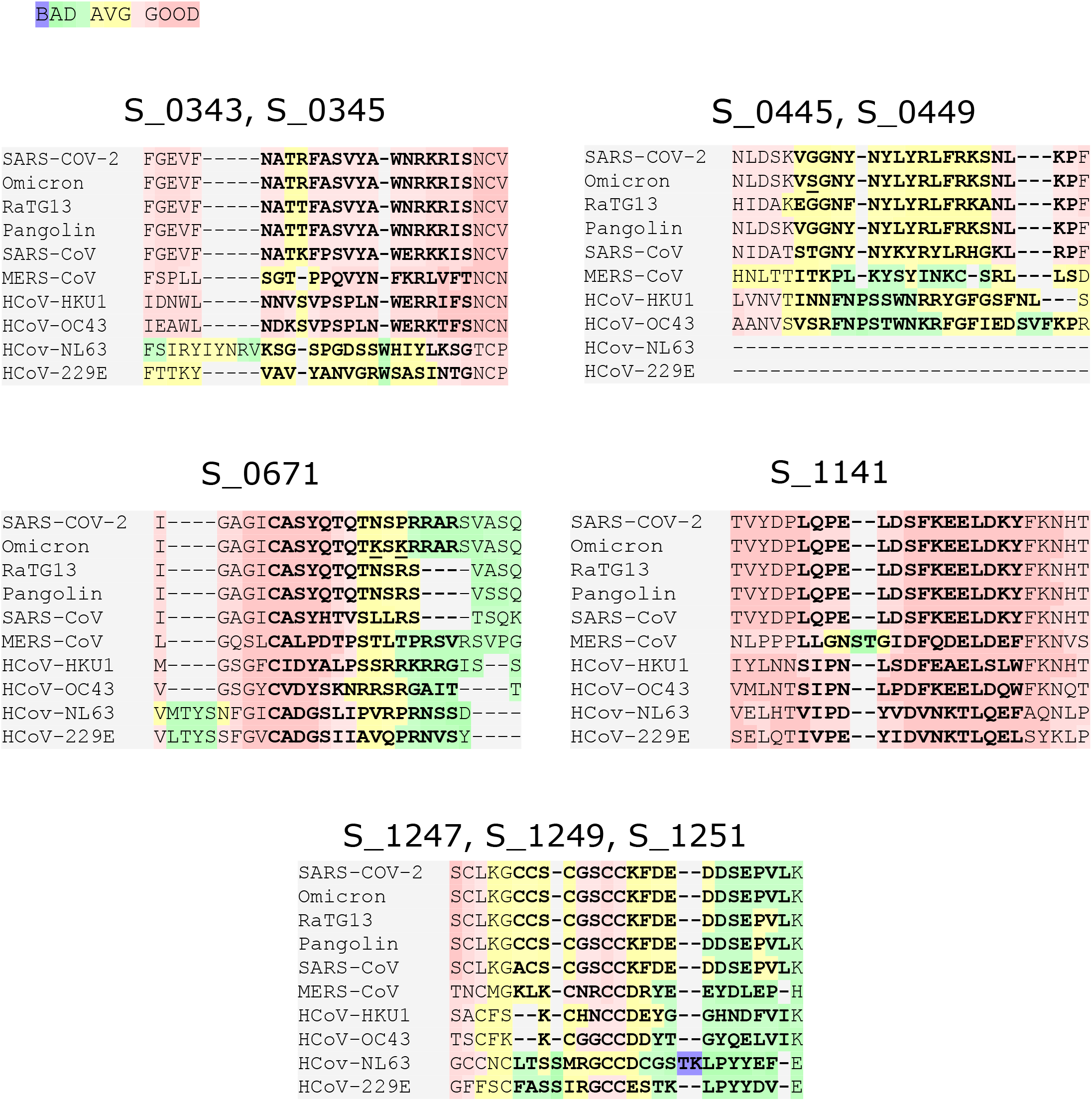
To determine conservation of the potential epitopes, the Wuhan SARS-CoV-2 sequence was aligned with the Omicron variant sequence, the other six human infected CoVs, and the closely related Pangolin and bat-infecting coronaviruses. Peptides IDs are displayed above their corresponding alignments and the color scale for the alignment can be found above.

### 3.2 Peptide Mutations of SARS-CoV-2 Variants Found in Epitopes of Interest

With the current prevalence of certain SARS-CoV-2 variants (21), it is important to assess whether their mutations are found in proximity to the identified peptides and whether these mutations have a role in increasing viral transmissibility. We discovered seven such variant mutation sites present in proximity to at least one of our peptides (Table 3) that are present in current prominent variants: Delta (B.1.617.2_India), Alpha (B.1.1.7_UK), and Omicron (B 1.1.529_South Africa).

**Table 3.** The seven recurring mutations of prevalent SARS-CoV-2 variants found within one of the 10 potential epitopes were identified to study their potential impact on antibody binding. Mutations are displayed in the leftmost column. Variants associated with each mutation are displayed in the middle column. Epitopes that contain the mutation are shown on the right.

| Mutation | Variant Found | Epitope(s) of Interest |
| --- | --- | --- |
| *L452R* | *Delta* | *S_0445, S_0449* |
| *G446D* | *Delta* | *S_0445* |
| *P681R* | *Delta* | *S_0671* |
| *P861H* | *Alpha* | *S_0671* |
| *G446S* | *Omicron BA.1* | *S_0445* |
| *N679K* | *Omicron* | *S_0671* |
| *P681K* | *Omicron* | *S_0671* |

The L452R and G446D mutations present in the Delta variant (22) were found within the region of the two overlapping identified epitopes S_0445 and S_0449. Although these mutations are not believed to directly affect the ACE2 binding activity of the virus, they are located in a peptide stretch that has been identified as an epitope for neutralizing antibodies and are thought to help inhibit neutralizing activity (23, 24). The G446S mutation found in the Omicron variant at the same position as the G446D mutation of the delta variant is also connected with escape from certain classes of neutralizing antibodies (25).

Another set of mutations, P681R, P681H, P681K, and N679K are all located within the peptide of interest S_0671. These mutations are commonly found in either Delta (P681R), Alpha (P681H), or Omicron (P681K and N679K) variants, within or in proximity to the polybasic insertion sequence PRRAR. This insertion sequence has been identified as the S1/S2 furin cleavage site that appears to be unique to SARS-CoV-2 among closely related CoVs (26, 27). Cleavage of this site gives RBD flexibility to change between an open and closed conformation before ACE2 Binding and is thought to be essential in conformational changes prior to viral host cell insertion (26, 27). The P681R mutation found in the Delta variant is thought to enhance furin cleavage function (28) and likely contributes to increased infectivity. It has been recently reported that the band of cleaved Omicron and Alpha were obviously weak despite the P681H mutation, which implies other mutations found in proximity to the furin cleavage site (T716I and N679K) could aid in the regulation of the proteolytic cleavage of S protein (29). Due to the significance of these mutations in the Delta and Omicron variants, the S_0671 peptide may be an epitope of particular interest for future studies. Assessing whether antibody binding to the mutated linear epitope is observed could help assess whether the mutation may also have a more direct role in the enhanced viral escape of the Delta and Omicron variants. The peptides found to be significant epitopes in our infected patient group, S_1141 and S_1247-S_1251, are located in the S2 subregion which is thought to be a highly conserved region among hCoVs (30). The peptides do not appear to contain any common mutations observed in the currently relevant variants (31).

It is interesting that none of the nine epitope sequences with significant IgG binding identified in the infected group were present in the N-terminal domain region of the spike protein. This was unexpected as the NTD region is thought to be a particularly important target of neutralizing antibodies (23, 24) and contains numerous mutations thought to help increase the infectivity of current variants (23, 25). These findings are corroborated by previous epitope mapping studies utilizing a proteome wide microarray that have identified few to no potential epitopes in this region (9, 11).

### 3.3 Antibody Binding Profile Comparison of Vaccinated and Infected Individuals

Our analysis of antibody immunoreactivity to the SARS-CoV-2 protein revealed evident differences between the antibody binding activity of the infected group and vaccinated individuals. Both groups showed overall significantly higher IgG binding activity to epitopes throughout the spike protein relative to the naïve samples. Despite similar levels of IgG binding activity between the two groups, distinct epitope binding patterns were only found in the infected group.

Previous studies have shown that vaccinated individuals produce a robust immune response that is comparable to natural infection and easily detectable with most methods (32, 33). We believe multiple factors were contributing to these unexpected results. Many of the acutely infected samples were collected two to three weeks after infection, when antibody production during infection is thought to peak (34). Corresponding symptom information for some infected patients was not available, so the number of infected patients with more severe symptoms is unknown. It has been found that severe symptoms correspond to higher antibody production (35) and more distinct antibody binding patterns (10, 11). If a large portion of these infected patients had severe symptoms, then their antibody binding would be expected to be higher. These variables limit comparisons between the infected and vaccinated group. From our findings, it would appear that the infected sample groups yielded more specific epitopes with significant IgG binding despite overall binding activity between the two groups being similar.

Although our results indicate that the vaccinated individuals did not have SARS-CoV-2 S protein epitopes that showed consistent binding, there were still some peptides that demonstrated the ability to potentially discriminate between vaccinated and naive individuals (S_1159 and S_1247-S_1251). These peptides did not meet the fold change criteria to be identified as significant in this study, so their potential sensitivity when incorporated into other assays is unknown. These peptides are well conserved among the hCoVs which means this enhanced binding may be non-specific. It is very possible that potential distinct epitopes with significant binding would appear in a larger sample pool of vaccinated individuals. Furthermore, it has been shown that previously infected individuals who receive a full vaccine treatment have extremely high antibody titers (36). The study of epitope binding among these individuals after their vaccination could lead to markers of vaccination to be more easily identified due to their higher antibody titers. Though no epitope markers were identified in the vaccinated group, this may be due to the relatively smaller sample size and low serum dilutions tested. An expanded study with a larger sample pool has high potential to yield epitopes with more distinct binding among vaccinated individuals.

The lack of significant IgA response within the spike region was somewhat surprising as previous studies had successfully identified epitopes within the spike protein utilizing peptide microarrays (11). Our ELISA data revealed relatively low IgA levels among some of the infected samples. A potential source of these lower IgA levels may be tied to a portion of the serum samples coming from asymptomatic and mildly symptomatic patients. IgA epitopes identified in previous peptide microarray studies (11) found that IgA epitopes were predominately identified among their severe symptom patient group. Since the quantity of overall IgA in serum is typically found to be significantly less than that of IgG, it is possible that levels of IgA present in some screened infected samples with milder symptoms were not high enough to discriminate from the naive samples. It has been observed that the amount of IgA antibodies reactive to the SARS-CoV-2 spike protein can vary dramatically among infected individuals, and significant decreases in IgA levels can be observed as early as four weeks after infection (36, 37).

Using SARS-CoV-2 peptide microarrays to perform IgA epitope profiling of sera may be difficult due to the limitations previously described without purification and concentration of IgA from samples. However, the potential for studying IgA epitopes of SARS-CoV-2 may still be viable with the use of saliva, which has a high content of IgA. Previous work using the peptide microarray to study the epitopes of other viral targets have shown the IgG antibody profiles of blood and saliva were similar (38). Salivary IgA has shown potential as a reliable biomarker for SARS-CoV-2 infection and a correlation between symptom severity and levels of salivary IgA have been previously demonstrated (39). Developing a method to treat microarrays with saliva may help to study IgA binding activity against SARS-CoV-2 epitopes. Previous attempts to directly treat microarrays with saliva are limited, but we have managed to obtain reproducible results screening processed saliva at low dilutions. Our ongoing study of IgG and IgA binding of saliva samples from previously infected individuals has demonstrated antibody binding to some epitopes identified in this report. These results are preliminary, with further analysis and validation required to draw any conclusions. Yet the screening of saliva samples may be helpful with efforts to perform IgA epitope profiling of the SARS-CoV-2 proteome and aid in the development of saliva-based diagnostics, immunity determinations, and therapeutics. This could also include detecting the immune response from the presence of the virus in the upper respiratory system within asymptomatic and pre-symptomatic individuals, possibly prior to detection by other methods.

Our comparison of the SARS-CoV-2 spike protein epitope profiles of vaccinated and infected individuals has led us to identify nine peptides with distinctly high IgG antibody reactivity in recently infected individuals. These nine peptides could potentially be implemented in diagnostic tools that could test for the presence of protective antibodies or active infection. Peptides with the potential to identify vaccine-only individuals have been identified, but further assessment of the use of said peptides to discriminate between vaccinated and naive individuals is necessary. The peptides may also serve to study the effect that mutations found in variants have on antibody binding. This information could be crucial as new variants continue to develop in the pandemic landscape.

