## Supplemental Table 4 for "Epitope Mapping of SARS-CoV-2 Spike Protein Reveals Distinct Antibody Binding Activity of Vaccinated and Infected Individuals"

| Sample ID | RBD | |
| --- | --- | --- |
| | IgA | IgG |
| 330 | 0.112825 | 2.033925 |
| 332 | 0.192575 | 3.040925 |
| 340 | 0.126225 | 3.771225 |
| 341 | 0.233125 | 3.082075 |
| 342 | 0.265175 | 3.041475 |
| 343 | 0.213975 | 3.681125 |
| 360 | 0.325575 | 3.774975 |
| 361 | 0.377075 | 3.879125 |
| 368 | 0.328075 | 3.632725 |
| 369 | 2.599825 | 3.731425 |
| 370 | 2.889825 | 3.710525 |
| 374 | 0.389475 | 3.738725 |
| 616 | 0.09065 | 0.7232 |
| 695 | 0.53395 | 3.9184 |
| 627 | 0.07895 | 0.0551 |
| 650 | 0.06985 | 0.0369 |
| 651 | 0.08775 | 0.0724 |
| 656 | 0.3944 | 3.8051 |
| 664 | 0.3287 | 3.8597 |
| 667 | 0.24535 | 1.72855 |
| 673 | 0.30715 | 3.8911 |
| 678 | 0.2326 | 1.45935 |
| 683 | 0.8013 | 3.88265 |
| 684 | 0.38825 | 3.9115 |
| 690 | 0.25585 | 3.8825 |
| 691 | 0.35025 | 3.89 |

| | | |
|---|---|---|
| 693 | 0.5371 | 3.9209 |
| 28823 | 0.088175 | 3.681125 |
| 28824 | 0.209725 | 3.774975 |
| 28825 | 0.077575 | 3.879125 |
| 28826 | 0.541775 | 3.731425 |
| V901C24 | 0.0743 | 1.81545 |
| V902C22 | 0.07075 | 3.72875 |
| V903C20 | 1.3552 | 3.8903 |
| V904C16 | 0.0859 | 2.965 |
| V907C17 | 0.9647 | 3.9126 |
| V951 | 0.5275 | 3.87465 |
| V957 | 0.70095 | 3.8883 |
| V960 | 0.41415 | 3.90295 |
| V963 | 0.4599 | 3.8985 |
| V964 | 0.50475 | 3.899 |
| V967 | 0.288 | 3.90915 |
| V975 | 0.5364 | 3.90465 |
| V977 | 0.60895 | 3.9036 |
| V982 | 0.41955 | 3.9101 |
| V987 | 0.7801 | 3.9058 |
| V989 | 1.6769 | 3.90105 |
| V900 | 0.76375 | 0.56705 |
| SN205 | 0.092525 | 0.0539 |
| SN206 | 0.192575 | 0.0365 |
| SN207 | 0.233125 | 0.0838 |
| SN210 | 0.265175 | 0.11465 |
| SN212 | 0.328075 | 0.059 |

| | | |
|---|---|---|
| V901A | 0.1828 | 0.2087 |
| V902A | 0.14885 | 0.0591 |
| V903A | 0.1612 | 0.1449 |
| V904A | 0.1792 | 0.33125 |
| V907A | 0.1518 | 0.46305 |
