## Supplemental Table 3 for "Epitope Mapping of SARS-CoV-2 Spike Protein Reveals Distinct Antibody Binding Activity of Vaccinated and Infected Individuals"

| ID | Age (years): | Sex | Race | Fever | Chills | Tiredness | Body Aches |
|---|---|---|---|---|---|---|---|
| PS616 | 59 | Female | No data | Severe | Severe | Severe | Moderate |
| PS650 | 24 | Female | No data | None | Mild | Moderate | Moderate |
| PS651 | 29 | Male | No data | None | Severe | Severe | Moderate |
| PS655 | 21 | Female | ɔanic or Lat | None | None | None | None |
| PS656 | 21 | Female | White | Moderate | Moderate | Moderate | Moderate |
| PS664 | 21 | Female | White | None | None | Mild | Mild |
| PS667 | 19 | Male | White | Moderate | Moderate | Moderate | Moderate |
| PS673 | 20 | Male | White | Mild | None | None | Mild |
| PS678 | 55 | Female | White | None | None | Mild | Mild |
| PS693 | 25 | Female | White | None | None | None | None |
| PS695 | 40 | Male | White | Moderate | Mild | Severe | Moderate |

| ID | Age (years): | Rash | Runny Nose | SOB | Cough | Pneumonia | Loss of Taste |
|---|---|---|---|---|---|---|---|
| PS616 | 59 | None | Moderate | Moderate | Severe | None | None |
| PS650 | 24 | None | Mild | Moderate | Moderate | None | Severe |
| PS651 | 29 | None | Severe | Moderate | Moderate | None | None |
| PS655 | 21 | None | None | None | None | None | None |
| PS656 | 21 | None | None | None | Mild | None | None |
| PS664 | 21 | None | Moderate | None | Mild | None | Mild |
| PS667 | 19 | None | Mild | Mild | Mild | None | None |
| PS673 | 20 | None | None | Moderate | Moderate | None | None |
| PS678 | 55 | None | None | Mild | None | None | Moderate |
| PS693 | 25 | None | None | None | None | None | None |
| PS695 | 40 | None | Mild | None | Mild | None | None |

| ID | Age (years): | Loss of Appetite | Sore Throat | Nausea | Diarrhea | Headache | Weakness |
|---|---|---|---|---|---|---|---|
| PS616 | 59 | Moderate | Moderate | None | Severe | Severe | Moderate |
| PS650 | 24 | Severe | Moderate | Mild | Mild | Severe | None |
| PS651 | 29 | None | Mild | Mild | Moderate | Severe | None |
| PS655 | 21 | None | None | None | None | None | None |
| PS656 | 21 | None | Moderate | None | None | Moderate | None |
| PS664 | 21 | Mild | Moderate | None | None | None | None |
| PS667 | 19 | Mild | None | None | None | Moderate | Mild |
| PS673 | 20 | None | None | None | None | Moderate | None |
| PS678 | 55 | None | None | None | None | Moderate | None |
| PS693 | 25 | None | None | None | None | None | None |
| PS695 | 40 | Mild | Mild | None | None | Severe | Mild |
