## Supplemental Table 2 for "Epitope Mapping of SARS-CoV-2 Spike Protein Reveals Distinct Antibody Binding Activity of Vaccinated and Infected Individuals"

| Sample ID | Sex | Age | Days Post Vaccine | Vaccine Received |
|---|---|---|---|---|
| *V901A** | *M* | *32* | *N/A* | *N/A* |
| *V902A** | *M* | *34* | *N/A* | *N/A* |
| *V903A** | *M* | *47* | *N/A* | *N/A* |
| *V904A** | *M* | *37* | *N/A* | *N/A* |
| *V907A** | *F* | *48* | *N/A* | *N/A* |
| *V901C24** | *M* | *32* | *24* | *Pfizer* |
| *V902C22** | *M* | *34* | *22* | *Moderna* |
| *V903C20** | *M* | *47* | *16* | *Moderna* |
| *V904C16** | *M* | *37* | *16* | *Pfizer* |
| *V907C17** | *F* | *48* | *17* | *Pfizer* |
| V951 | M | 47 | 19 | Moderna |
| V957 | F | 51 | 17 | Pfizer |
| V960 | F | 25 | 28 | Pfizer |
| V963 | F | 26 | 19 | Moderna |
| V964 | M | 58 | 9 | Pfizer |
| V967 | M | 66 | 17 | Moderna |
| V975 | M | 67 | 44 | Moderna |
| V977 | F | 36 | 10 | Pfizer |
| V982 | M | 22 | 21 | Pfizer |
| V987 | M | 62 | 20 | Moderna |
| V989 | M | 24 | 6 | Pfizer |
| V900 | M | 65 | 108 | Moderna |
| SN205 | F | 22 | N/A | N/A |
| SN206 | M | 18 | N/A | N/A |
| SN207 | M | 47 | N/A | N/A |
| SN210 | M | 28 | N/A | N/A |
| SN212 | F | 39 | N/A | N/A |
