## Supplemental Table 1 for "Epitope Mapping of SARS-CoV-2 Spike Protein Reveals Distinct Antibody Binding Activity of Vaccinated and Infected Individuals"

| Sample ID | Sex | Age | Day of Test | Date of Sample Collection | Symptom Information |
|---|---|---|---|---|---|
| 330 | F | 60 | 4/2/2020 | 5/3/2020 | Unavailable |
| 332 | F | 56 | 4/1/2020 | 5/3/2020 | Unavailable |
| 340 | F | 73 | 4/2/2020 | 5/3/2020 | Unavailable |
| 341 | F | 65 | 4/1/2020 | 5/3/2020 | Unavailable |
| 342 | M | 83 | 4/2/2020 | 5/3/2020 | Unavailable |
| 343 | M | 71 | 4/3/2020 | 5/3/2020 | Unavailable |
| 360 | M | 67 | 4/17/2020 | 5/17/2020 | Unavailable |
| 361 | F | 68 | 4/17/2020 | 5/17/2020 | Unavailable |
| 368 | F | 75 | 4/17/2020 | 5/17/2020 | Unavailable |
| 370 | M | 64 | 4/17/2020 | 5/17/2020 | Unavailable |
| 374 | F | 76 | 4/17/2020 | 5/17/2020 | Unavailable |
| 616 | F | 59 | 8/5/2020 | 9/9/2020 | Severe |
| 627 | F | 22 | 9/14/2020 | 9/23/2020 | Severe |
| 650 | F | 24 | 10/14/2020 | 10/21/2020 | Moderate |
| 651 | M | 29 | 10/15/2020 | 10/21/2020 | Moderate |
| 656 | F | 21 | 11/2/2020 | 11/11/2020 | Moderate |
| 664 | F | 21 | 11/7/2020 | 11/11/2020 | Mild |
| 667 | M | 19 | 11/16/2020 | 11/19/2020 | Moderate |
| 673 | M | 20 | 11/11/2020 | 12/4/2020 | Mild |
| 678 | F | 55 | 12/9/2020 | 12/11/2020 | Mild |
| 683 | F | 44 | 12/21/2020 | 1/6/2021 | Moderate |
| 684 | M | 34 | 12/18/2020 | 1/6/2021 | Unavailable |
| 690 | F | 31 | 1/2/2021 | 1/20/2021 | Unavailable |
| 691 | M | 31 | 1/8/2021 | 1/20/2021 | Unavailable |
| 693 | F | 25 | 1/10/2021 | 1/27/2021 | Unavailable |
| 695 | M | 40 | 1/2/2021 | 2/3/2021 | Moderate |
| 28823 | M | 44 | 11/12/2020 | 11/12/2020 | Unavailable |

| | | | | | |
|---|---|---|---|---|---|
| 28824 | M | 60 | 12/6/2020 | 12/6/2020 | Unavailable |
| 28825 | F | 28 | 12/31/2020 | 12/31/2020 | Unavailable |
| 28826 | F | 66 | 11/11/2020 | 11/11/2020 | Unavailable |
